# Autophagy enhances memory erasure through synaptic destabilization

**DOI:** 10.1101/237636

**Authors:** Mohammad Shehata, Kareem Abdou, Kiriko Choko, Mina Matsuo, Hirofumi Nishizono, Kaoru Inokuchi

## Abstract

There is substantial interest in memory reconsolidation as a target for the treatment of anxiety disorders such as post-traumatic stress disorder (PTSD). However, its applicability is restricted by reconsolidation-resistant conditions that constrain the initial memory destabilization. In this study, we investigated whether the induction of synaptic protein degradation through autophagy modulation, a major protein degradation pathway, can enhance memory destabilization upon retrieval and whether it can be utilized to overcome these conditions. Here, using male mice in an auditory fear reconsolidation model, we showed that autophagy contributes to memory destabilization and its induction can be utilized to enhance erasure of a reconsolidation-resistant auditory fear memory that depended on α-amino-3-hydroxy-5-methyl4-isoxazolepropionic acid receptor (AMPAR) endocytosis. Using male mice in a contextual fear reconsolidation model, autophagy induction in the amygdala or in the hippocampus enhanced fear or contextual memory destabilization, respectively. The latter correlated with AMPAR degradation in the spines of the contextual memory-ensemble cells. Using male rats in an *in vivo* long-term potentiation reconsolidation model, autophagy induction enhanced synaptic destabilization in an N-methyl-D-aspartate receptor-dependent manner. These data indicate that induction of synaptic protein degradation can enhance both synaptic and memory destabilization upon reactivation and that autophagy inducers have the potential to be used as a therapeutic tool in the treatment of anxiety disorders.

**Significance Statement:** It has been reported that inhibiting synaptic protein degradation prevents memory destabilization. However, whether the reverse relation is true and whether it can be utilized to enhance memory destabilization is still unknown. Here we addressed this question on the behavioral, molecular and synaptic levels, and showed that induction of autophagy, a major protein degradation pathway, can enhance memory and synaptic destabilization upon reactivation. We also show that autophagy induction can be utilized to overcome a reconsolidation-resistant memory, suggesting autophagy inducers as a potential therapeutic tool in the treatment of anxiety disorders.

## Introduction

Retrieval of long-term memories (LTM) can induce a destabilization process that returns them into a labile state, which is followed by a protein synthesis-dependent reconsolidation process that serves to strengthen or update the original memories (Nader et al., 2000; Besnard et al., 2012; Finnie and Nader, 2012; Inaba et al., 2015; Lee et al., 2017). Blocking reconsolidation has been suggested as a tool to weaken traumatic memories in anxiety disorders such as post-traumatic stress disorder (PTSD). However, the initial destabilization step is challenging when memories are formed under extremely stressful conditions, and it would require pharmacological assistance (Tronson and Taylor, 2007; Pitman, 2011; Besnard et al., 2012). It has been reported that inhibition of synaptic protein degradation, through blocking the ubiquitin-proteasome system, prevents memory destabilization (Lee et al., 2008). However, whether induction of synaptic protein degradation can be utilized to enhance memory destabilization is yet to be tested.

Macro-autophagy, hereafter referred to as autophagy, is a major protein degradation pathway where a newly synthesized isolation membrane sequesters a small portion of the cytoplasm to form a multilamellar vesicle called an autophagosome. To degrade the entrapped contents, autophagosomes fuse into the endosome-lysosome system (Mizushima and Komatsu, 2011; Yamamoto and Yue, 2014). The process of autophagosome synthesis is orchestrated by molecular machinery consisting of the autophagy-related genes (Atg) found in yeast, and their mammalian homologs (Mizushima et al., 2011; Ohsumi, 2014). In the brain, autophagy plays an important role in neurodegenerative diseases (Yamamoto and Yue, 2014) and is essential for the development of a healthy brain (Hara et al., 2006; Komatsu et al., 2006; Liang et al., 2010). It has been suggested that neurons may have adapted autophagy to suit their complex needs including contribution to synaptic function (Bingol and Sheng, 2011; Mizushima and Komatsu, 2011; Shehata and Inokuchi, 2014; Yamamoto and Yue, 2014). In line with this idea, autophagosomes are found not only in the neuron’s soma and axons but also in the dendrites (Hollenbeck, 1993; Shehata et al., 2012), Also, autophagy contributes to the degradation of the endocytosed γ-aminobutyric acid receptors (GABAR) in *Caenorhabditis elegans* and of the α-amino-3-hydroxy-5-methyl4-isoxazolepropionic acid receptors (AMPAR) upon chemical longterm depression (LTD) in cultured neurons (Rowland et al., 2006; Shehata et al., 2012). Both GABAR and AMPAR play pivotal roles in the synaptic plasticity models of LTD and long-term potentiation (LTP), which are causally correlated with memory (Kessels and Malinow, 2009; Squire and Kandel, 2009; Nabavi et al., 2014). Moreover, the regulation of autophagy intersects with protein synthesis regulation at the mammalian target of rapamycin (mTOR) and the phosphatidylinositol-3-monophosphate kinase (PI3K) and by careful consideration of the discrepancy in the effects of the mTOR and PI3K modulators on memory processes, autophagy is suggested to play a role in memory reconsolidation (Chen et al., 2005; Gafford et al., 2011; Shehata and Inokuchi, 2014). In the present study, we tested the hypothesis that autophagy could play a role in synaptic and memory destabilization and therefore, the induction of autophagic protein degradation can be utilized to enhance erasure of reconsolidation-resistant fear memories.

## Materials & Methods

### Animals

All procedures involving the use of animals were conducted in compliance with the guidelines of the National Institutes of Health (NIH) and were approved by the Animal Care and Use Committee of the University of Toyama, Japan. Eight-week-old male Wistar ST rats were purchased for electrophysiological experiments, and 8-week-old male C57BL/6J mice were purchased for behavioral experiments. The c-fos-tTA mice were purchased from the Mutant Mouse Regional Resource Center (stock number: 031756-MU). The progeny for the c-fos-tTA line was generated using *in vitro* fertilization of eggs from C57BL/6J mice, as described previously (Ohkawa et al., 2015). All animals were purchased from Sankyo Labo Service Co. Ltd. (Tokyo, Japan). Rats and mice were maintained in separate rooms on a 12 h light-dark cycle at 24 ± 3°C and 55 ± 5% humidity. They were given food and water ad libitum and housed with littermates until surgery.

### Drugs and peptides

Anisomycin (Sigma Aldrich Japan Co., Tokyo, Japan) was dissolved in a minimum quantity of HCl, diluted with phosphate buffered saline (PBS), and adjusted to pH 7.4 with NaOH (Ani). Ifenprodil tartrate (Sigma Aldrich Co., Japan) and trifluoperazine dihydrochloride (Sigma Aldrich Japan Co.) were dissolved in PBS. Spautin-1 (Sigma Aldrich Co., Japan) was dissolved in DMSO and diluted with equal volume of saline. The retro-inverso Tat-beclin 1 peptide D-amino acid sequence (RRRQRRKKRGYGGTGFEGDHWIEFTANFVNT; synthesized by GenScript through Funakoshi Co., Ltd., Tokyo, Japan) was dissolved in either PBS (tBC) or anisomycin solution (Ani+tBC). The control D-Tat peptide D-amino acid sequence (YGRKKRRQRRR; EMC microcollections, Tübingen, Germany) was dissolved in PBS (D-Tat). The Tat-GluA2_3Y_ peptide L-amino acid sequence (YGRKKRRQRRRYKEGYNVYG, AnaSpec, Fremont, CA) and its control Tat-GluA2_3A_ peptide L-amino acid sequence (YGRKKRRQRRRAKEGANVAG; AnaSpec), were both dissolved in PBS (GluA2_3Y_ or GluA2_3A_, respectively). All peptides were aliquoted into single experiment volumes and stored at -80°C.

### Stereotactic surgery and drug infusion in mice

Mice were 8–10 weeks old at the time of surgery. They were anesthetized with isoflurane, given an intraperitoneal injection of pentobarbital solution (80 mg/kg of body weight), and then placed in a stereotactic apparatus (Narishige, Tokyo, Japan). Mice were then bilaterally implanted with a stainless guide cannula (PlasticsOne, Roanoke, VA, USA). For targeting the CA1, the guide cannula was positioned 1.8 mm posterior, 1.55 mm lateral, and 1.5 (C57BL/6J) or 1.0 mm (c-fos-tTA mice) ventral to the bregma. For targeting the BLA, the guide cannula was positioned 1.5 mm posterior, 3.3 mm lateral, and 3.4 mm ventral to the bregma. For targeting the LA, the guide cannula was positioned 1.7 mm posterior, 3.4 mm lateral, and 2.6 mm ventral to the bregma. After surgery, a cap or dummy cannula (PlasticsOne) was inserted into the guide cannula, and mice were allowed to recover for at least 7 days in individual home cages before the experiment. Mice in the NoFS condition were not cannulated. All drug infusions were done under isoflurane anesthesia, using an injection cannula with a 0.25 mm internal diameter (PlasticsOne), and extending beyond the end of the guide cannula by 0.5 mm for the CA1, or by 1.5 mm for the BLA and LA. The drug infusion rate was 0.2 μl/minute for the CA1 in C57BL/6J mice, or 0.1 μl/minute for the CA1 in c-fos-tTA mice, and the BLA and LA. Following drug infusion, the injection cannula was left in place for 2 minutes to allow for drug diffusion. For the reconsolidation experiments, immediately after retrieval 1 μl of drug solution was injected into the CA1 in C57BL/6J mice, or 0.5 μl was injected into the CA1 in c-fos-tTA mice, the BLA and LA. In all of these reconsolidation experiments, 1 μl of drug solution contained either PBS, 125 μg of Ani, 20 μg of tBC, or 125 μg Ani + 20 μg tBC. For autophagy inhibition, 0.5 μl of a solution containing 8.3 μg Spautin-1 or vehicle was injected into LA. For blocking AMPA receptor endocytosis, 0.5 μl of solution containing 20 ng of GluA2_3Y_ or GluA2_3A_ was injected into the LA.

### Lysate preparation and immunoblot analysis

Drugs were infused into the CA1 or amygdala of one hemisphere of the C57BL/6J mice, as described above. Four hours later, their brains were removed and cut into 1-mm slices, placed on ice, and the hippocampus or amygdala from each hemisphere was dissected under a binocular microscope, rapidly frozen on dry ice, and stored at -80°C. Samples were then sonicated in RIPA buffer (50 mM TrisHCl, pH 7.5, 150 mM NaCl, 2 mM EDTA, 1% NP-40, 0.5% sodium deoxycholate, 0.1% SDS, and 50 mM NaF) containing a protease inhibitor mixture (cOmplete ULTRA tablets, Roche Diagnostics GMBH, Mannheim, Germany) and a phosphatase inhibitor mixture (PhosSTOP tablets, Roche Diagnostics GMBH). Samples were then centrifuged at 14000 rpm for 15 minutes at 4°C, and supernatants were stored at -30°C until use. Measurement of protein concentration, immunoblotting for LC3 detection (ab48394; Abcam, Tokyo, Japan), visualization and quantitation were performed as previously described (Shehata et al., 2012).

### Contextual fear conditioning (CFC)

All behavioral sessions were conducted during the light cycle, in a dedicated soundproof behavioral room (Yamaha Co., Shizuoka, Japan), described here as Room A. The CS context was a square chamber (Chamber A) with a plexiglass front, off-white side- and back-walls (length 175 × width 165 × height 300 mm) and a floor consisting of stainless steel rods connected to an electric shock generator. The distinct context was a circular chamber (Chamber B) with opaque reddish walls (diameter 235 mm × height 225 mm) and a smooth gray floor. One day before the experiment, mice were left undisturbed on a waiting rack for 2 h for habituation purposes. On the day of the experiment, mice were left undisturbed on the waiting rack for at least 30 minutes before and after each session, and during the experiment. In each session, one mouse in its home cage was moved into Room A. During the conditioning or reconditioning sessions, mice were placed in Chamber A and allowed to explore for 148 sec, before receiving one footshock (2 sec, 0.4 mA). They then remained for 30 sec, before being moved back to their home cages and returned to the waiting rack. During the retrieval session (T1), mice were placed back into Chamber A for 3 minutes, then immediately subjected to isoflurane anesthesia and drug infusion. Mice in the NoFS condition were manipulated identically, with the exceptions that the shock generator was turned off. During the test sessions, mice were placed back into Chamber A (T2 and T4) for 5 minutes, and 1 h later into Chamber B (T3) for 5 minutes. Mice remained on the waiting rack during the 1 h interval. In all behavioral sessions, chambers were cleaned with 70% ethanol and water between each mouse, and kept odorless to the experimenter.

### Auditory fear conditioning (AFC)

Different chambers were used for each auditory fear conditioning session. Context exploration and conditioning were performed in Chamber A. Retrieval sessions were performed in a circular chamber (Chamber C) with opaque black walls (diameter 215 mm × height 340 mm) and a smooth gray floor. Test sessions were performed in a circular chamber (Chamber D) with opaque reddish walls (diameter 235 mm × height 310 mm) and a smooth gray floor. After recovery from surgery, a maximum of six mice were moved with their home cages on racks in the maintenance room to a soundproof (Yamaha Co.) waiting room (Room B). Mice were left undisturbed for at least 15 minutes before and after each session and during the experiment. In each session, one mouse in its home cage was moved into Room A. During the context exploration sessions, mice were placed in Chamber A and allowed to explore for 5 minutes per day for 2 days. During the conditioning sessions, mice were placed in Chamber A for 2 minutes, and then received one or three tones (30 sec, 65 dB, 7 kHz), co-terminating with a shock (2 sec, 0.4 mA), with an interval of 30 sec. After the last shock, mice remained for 30 sec, and were then returned to their home cages and to Room B. During the retrieval sessions, mice were placed into Chamber C for 2 minutes before receiving a tone (30 sec, 65 dB, 7 kHz), then 30 sec later, mice were subjected to isoflurane anesthesia and drug infusion before being returned to Room B. For autophagy inhibition or blocking of AMPA receptor endocytosis, mice were subjected to isoflurane anesthesia and drug infusion 75 minutes prior to the retrieval sessions. During test sessions, mice were placed in Chamber D for 2 minutes, and then received a tone (30 sec, 65 dB, 7 kHz).

### Behavioral analysis

All experiments were conducted using a video tracking system (Muromachi Kikai, Tokyo, Japan) to measure the freezing behavior of the animals. Freezing was defined as a complete absence of movement, except for respiration. We started scoring the duration of the FR after 1 sec of sustained freezing behavior. All behavioral sessions were screen recorded using Bandicam software (Bandisoft, Seoul, Korea). Occupancy plots representing the maximum occupancy of the mouse center in the defined context space during each session were generated by analyzing the screen recorded movies using ANY-maze software (Stoelting Co., Wood Dale, IL, USA). Mice were assessed as completely amnesic when they: (1) showed at least a 50% decrease in freezing level after drug infusion compared with the level before treatment, and (2) showed a freezing level in the CS or distinct contexts within the 95% confidence interval of the freezing level of the NoFS condition (used as a reference for normal mouse behavior). Animals were excluded from behavioral analysis when showing abnormal behavior after surgery or the cannula was misplaced in position.

### Plasmid construction, lentivirus preparation, and infection

For plasmid construction, mCherry (Clontech, Palo Alto, CA, USA) was amplified by PCR using the following primers, sense: gggggatccgccaccatggtgagcaagggcgaggagg; antisense: ggggtcgaccccgggctacttgtacagctcgtcc. The resulting fragment was then used to replace the EYFP fragment at the BamHI-Sall sites in pBS-TRE3G-EYFP to produce the pBS-TRE3G-mCherry plasmid. The pBS-TRE3G-EYFP plasmid is a pBluescript II SK+ plasmid (Stratagene, La Jolla, CA, USA) containing the third generation tTA-responsive TRE3G promoter sequence, derived from pTRE3G-IRES (Clontech, 631161) fused to EYFP. Finally, the TRE3G-mCherry fragment was subcloned into the STB plasmid using the SpeI/XbaI-XmaI sites to produce the pLenti-TRE3G-mCherry plasmid, which was used for the lentivirus preparation as previously described (Ohkawa et al., 2015). The viral titer was approximately 5 × 10^9^ IU/ml. Virus infection into CA1 of the c-fos-tTA mice (18–20 weeks old) was performed during the surgery for drug cannula fixation. Lentivirus (0.5 μl/site) was introduced through an injection cannula inserted into the guide cannula and left protruding by 0.5 mm. The injection rate was 0.1 μl/minute, and the cannula was left in place for 20 minutes after the end of the injection, before being slowly withdrawn.

### Labeling of the memory-ensemble cells

Labeling of the memory-ensemble cells associated with contextual fear was performed in a similar manner to the experiments on CFC. The experiment was performed on lentivirus-injected c-fos-tTA mice, maintained since weaning on food containing 40 mg/kg doxycycline. Two weeks after lentivirus infection, mice were subjected to the waiting rack for 2 h for habituation purposes. One day later doxycycline was removed and mice were maintained on normal food. Two days after doxycycline removal, mice were subjected to a CFC session as mentioned above. Six hours later, the feed for the mice was changed to food containing 1000 mg/kg doxycycline. The retrieval session and drug infusion were performed as mentioned above. One day after drug infusion, the mice were deeply anesthetized with an overdose of pentobarbital solution, and perfused transcardially with PBS (pH 7.4) followed by 4% paraformaldehyde in PBS (PFA). The brains were removed, further post-fixed by immersion in PFA for 16–24 h at 4°C, equilibrated in 30% sucrose in PBS for 36–48 h at 4°C, and then stored at −80°C.

### Immunohistochemistry

Double labeling primary antibodies from the same host species (rabbit) were used for GluA1 and mCherry staining. Several incisions where made to label the right side of the brains, and they were then cut into 50 μm coronal sections using a cryostat and transferred to 12-well cell culture plates (Corning, NY, USA) containing PBS. After washing with PBS, the floating sections were treated with blocking buffer (5% normal donkey serum; S30, Chemicon by EMD Millipore, Billerica, MA USA) in 0.3% Triton X-100-PBS (TPBS) at room temperature (RT) for 1 h. They were then treated with anti-GluA1 antibody (1:500; AB1504; EMD Millipore) in blocking buffer at 4°C for 36-40 h. After three 10-minute washes with 0.1% PBST (the procedure for further mentions of washing in this paragraph), sections were incubated with donkey anti-rabbit IgG-AlexaFluor 488 secondary antibody in blocking buffer (A21206, Molecular Probes, Invitrogen, Carlsbad, CA, USA) at RT for 4 h. After washing, sections were incubated with 5% normal rabbit serum (Jackson ImmunoResearch Inc., West Grove, PA, USA) in 0.3% TPBS at RT for 1 h. Following washing, sections were incubated with 4% Fab Fragment Donkey Anti-Rabbit IgG (Jackson ImmunoResearch) in 0.3% TPBS at RT for 2 h. Sections were then washed and treated with anti-mCherry antibody (1:500; 632496; Clontech) in blocking buffer at 4°C for 36-40 h. After washing, sections were incubated with donkey anti-rabbit IgG-AlexaFluor546 secondary antibody in blocking buffer (A10040, Molecular Probes) at RT for 4 h. Sections were then washed and treated with DAPI (1 μg/ml, Roche Diagnostics, 10236276001), then washed three times with PBS. Sections were then mounted on glass slides with ProLong Gold antifade reagent (Invitrogen).

### Confocal microscopy and analysis of puncta

Images were acquired using a Zeiss LSM 780 confocal microscope (Carl Zeiss, Jena, Germany). First, a Plan-Apochromat 5× objective lens was used to check for the treatment side, and then low magnification images of the CA1 radiatum were acquired for each selected hemisphere using a Plan-Apochromat 20× objective lens. High magnification images for dendrites and spines were acquired using a Plan-Apochromat 63×/1.4 oil DIC objective lens. All acquisition parameters were kept constant within each magnification. To detect GluA1 puncta and the mCherry-labeled dendrites and spines, high resolution (4096 × 4096) images were acquired by collecting z-stacks (5 slices at 0.6 μm thickness, and 0.3 μm interval). After performing a digital zoom (7×), maximum intensity projection images were created with ZEN 2.1 Black (Carl Zeiss) and further processed with Gamma correction at γ = 1.5 (for mCherry-labeled spines and Total GluA1 puncta) or γ = 5 (for GluA1 strong signal). ImageJ software (NIH, Bethesda, Maryland, USA) was used to apply a constant threshold to the green channel to create binary images for both Total GluA1 puncta (GluA(Tot)^+^) and GluA1 strong signal (GluA1(Str)^+^). Both puncta were automatically counted using the Analyze particles function with a particle size of > 50 pixel^2^ for GluA(Tot)^+^ or > 100 pixel^2^ for GluA1(Str)^+^, and a circularity of 0.2−1.0. Any fused puncta were manually separated before automatic counting. Overlaps between the GluA1(Str)^+^ puncta and mCherry^+^ spines were manually counted, guided by the green and red color thresholding in ImageJ. The mCherry^+^ only spines did not overlap with any GluA1(Tot)^+^ puncta. Three hemispheres were analyzed for each of the PBS or Ani+tBC treatments from four mice. For each hemisphere, data from four analyzed maximum intensity projection images were averaged.

### Stereotactic surgery and drug infusion in rats

Previously described surgical procedures were used with some modifications (Okubo-Suzuki et al., 2016). Rats were 8−10 weeks old at the time of surgery. In brief, a bipolar stimulating electrode and a monopolar recording electrode, both made of tungsten wire, were stereotaxically positioned to stimulate the perforant pathway (angular bundle), while recording in the dentate gyrus. The stimulating electrode was positioned 7.5 ± 0.3 mm posterior, 4.4 ± 0.3 mm lateral, and 4.7 ± 0.3 mm ventral to the bregma. The recording electrode was positioned ipsilaterally 4.0 ± 0.3 mm posterior, 2.5 ± 0.3 mm lateral and 3.8 ± 0.3 mm ventral to the bregma. For intracerebroventricular (ICV) infusion, a stainless-steel guide cannula (Eicom, Tokyo, Japan) was positioned ipsilaterally 0.7 ± 0.3 mm posterior, 1.6 ± 0.3 mm lateral, and 4.0 mm ventral to the bregma. After surgery, a dummy cannula (Eicom), which extended 1.0 mm beyond the end of the guide cannula, was inserted into the guide cannula. Rats were allowed to recover for at least 10 days in individual home cages before the experiment. ICV drug infusion was performed on unanesthetized freely moving rats, using an injection cannula (Eicom) that extended 0.5 mm beyond the end of the guide cannula, with an infusion rate of 1 μl/minute. Following drug infusion, the injection cannula was left in place for 5 minutes to allow for drug diffusion.

### In vivo electrophysiology on freely moving rats

The LTP experiments were modified from those previously described (Okubo-Suzuki et al., 2016). After recovery from surgery, the input/output curves were determined as a function of current intensity (0.1−1.0 mA), and the intensity of the stimulus current required to elicit the maximum fEPSP slope (MAX) was determined for each animal. The stimulus current intensity was set to elicit 50% of MAX. Three days later, 400 Hz or 8 Hz stimulations were induced. The 400 Hz stimulation used for LTP induction consisted of 10 trains with 1 minute inter-train intervals, with each train consisting of five bursts of 10 pulses at 400 Hz, delivered at 1 s interburst intervals, giving a total of 500 pulses. The later 8 Hz stimulation, which was performed as a reactivation stimulation, consisted of 128 pulses at 8 Hz. The fEPSP slope was monitored by delivering test pulses at 0.05 Hz for 15 minutes before (PreStim), and 5 minutes after (PostStim) stimulation, over the following 4 days. For testing the dependency of the stimulation on protein synthesis, 5 μl PBS or 5 μl of a solution containing 400 μg of Ani, were infused directly after the PostStim recording. For LTP reconsolidation experiments, LTP (400 Hz stimulation) was induced 3 days after MAX, and 1 day later, the 8 Hz reactivation stimulation was performed. As mentioned above, the fEPSP slope was monitored both before (PreLTP) and 5 minutes after LTP induction (PostLTP), and before (PreReact) and 5 minutes after (PostReact) the 8 Hz reactivation stimulation. Immediately after PostReact recording, rats received a 5 μl IVC drug infusion, containing either PBS, 400 μg of Ani, 100 μg of tBC, or 400 μg of Ani + 100 μg of tBC, as described above. For inhibition of NMDAR-2B, 5 μl of a solution containing 5 μg of ifenprodil tartrate was IVC infused immediately before the PreReact recording. The fEPSP slope was monitored over the following 3 days. Rats were excluded when showing abnormal behavior after surgery, LTP was not induced from the first trial, or the cannula or the electrodes were misplaced in position.

### Experimental Design and Statistical Analysis

In figure legends, n refers to the number of animals per treatment condition unless otherwise indicated. All experiments were performed at least three times with lots of 3-6 animals each. Treatments were counterbalanced for each lot. Animals were blindly and randomly allocated for each treatment condition. Statistical analysis was performed using Prism 6.01 or InStat 3.1 (GraphPad Software, San Diego, CA, USA). Data from two conditions were compared using two-tailed unpaired Student t tests. Multiple-condition comparisons were assessed using ANOVA with post hoc tests as described the results section. *P* values were considered significant if less than 0.05. Quantitative data are presented as mean ± SEM.

## Results

### Autophagy contributes to fear memory destabilization

To modulate autophagy activity within the time window of reconsolidation, we pharmacologically targeted the Beclin1 protein, which is part of the Beclin1-Atg14L-Vps34 lipid kinase complex that is involved in the autophagosome synthesis. This will specifically modulate autophagy activity without affecting endocytosis, mTOR, or PI3K activity (Vanhaesebroeck et al., 2010; Liu et al., 2011; Shoji-Kawata et al., 2013; Marsh and Debnath, 2015; De Leo et al., 2016). For autophagy induction, we used the cell-permeable tat-beclin1 peptide (tBC), which is composed of the human immunodeficiency virus-1 transduction domain attached to the necessary and sufficient peptide sequence of the beclin1 protein (Shoji-Kawata et al., 2013). The tBC peptide induces autophagy in the brains of mice neonates when systemically injected (Shoji-Kawata et al., 2013), and induces autophagy in the amygdala of adult mice when directly infused as monitored through the conversion of the light chain protein 3 (LC3), an autophagosome-specific marker, from its inactive form (LC3-I) to the lipidated active form (LC3-II). For autophagy inhibition, we used Spautin-1 (Spautin), which promotes the degradation of the Beclin1-Atg14L-Vps34 complex through inhibiting the ubiquitin-specific peptidases that target the beclin1 subunit of the complex (Liu et al., 2011). Infusion of spautin into the amygdala inhibited both the basal and the tBC-induced autophagic activity (one-way ANOVA, LC3-II/LC3-I: *P* = 0.007 and Total LC3: *P* = 0.505; Tukey’s post-hoc test; Fig. 1*A*,*B*).

**Figure 1.**
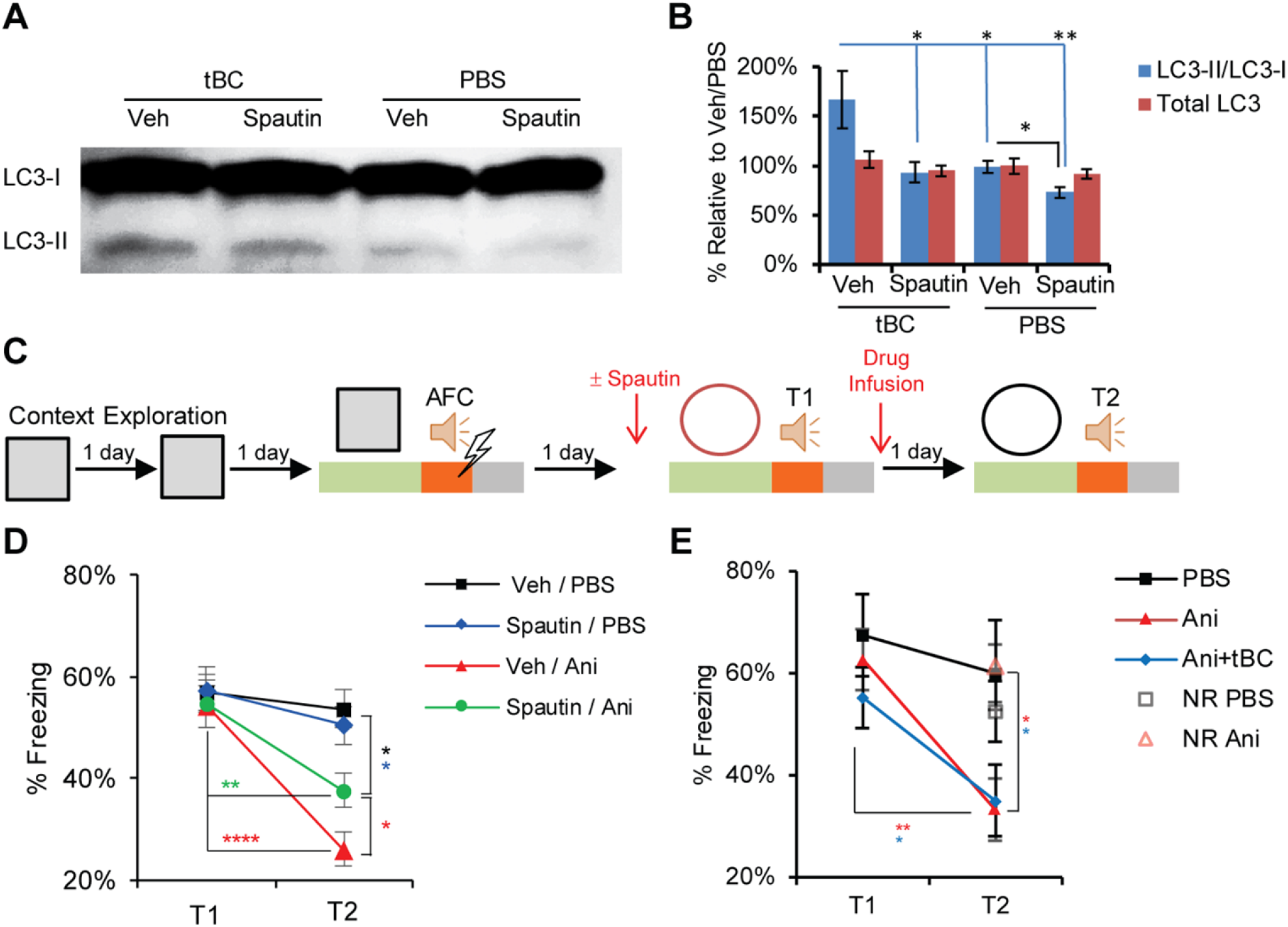
Autophagy contributes to fear memory destabilization. ***A***, Representative LC3 immunoblot from mouse amygdala lysates showing autophagy induction by Tat-beclin 1 (tBC) and inhibition by Spautin-1 (Spautin). The lipidated active form (LC3-II) is an autophagosome-specific marker. ***B***, Quantitation of the immunoblot signal intensity represented as percentage relative to a Veh/PBS sample (n = 4 mice/condition). ***C***, Design for the one tone-footshock pair auditory fear conditioning (1FS-AFC) experiments. ***D***, Average percentage freezing during tone at T1 and T2 showing that blocking autophagy significantly decreased the amnesic effect of Ani (n = 10-11 mice/condition). ***E***, Average percentage freezing during tone at T1 and T2 showing that, when the amnesic effect of anisomycin (Ani) was complete, autophagy induction did not have any further effect. No reactivation (NR) control showed no amnesic effect. Note: no injections were done before T1. (n = 7-9 mice/condition). Error bars represent mean ± SEM; * *P* < 0.05; ** *P* < 0.01; **** *P* < 0.0001. PBS: phosphate buffered saline; Veh: Vehicle.

To examine the effect of autophagy modulation on memory destabilization, we employed a reconsolidation model of fear conditioning. Fear conditioning is an associative learning paradigm, in which animals learn to associate a specific auditory cue (auditory fear conditioning, AFC) or context (contextual fear conditioning, CFC), which is a conditional stimulus (CS), with a foot shock, an unconditional stimulus (US). When animals are subjected to the CS, they recall the fear memory, resulting in a freezing response.

When a one tone-footshock pair (1FS-AFC) was used for conditioning in the AFC paradigm, anisomycin (Ani) infusion into the lateral amygdala (LA) after tone retrieval led to a significant decrease in the tone-elicited freezing response compared with the vehicle-infused condition (Fig. 1*C-E*), in agreement with previous reports (Nader et al., 2000; Suzuki et al., 2004; Mamiya et al., 2009). Ani produced a retrieval-specific retrograde amnesia as Ani administration without the retrieval session had no effect on tone fear memory (Fig. 1*E*). Inhibiting autophagy through Spautin infusion into the LA before retrieval partially blocked the Ani amnesic effect, indicating that autophagy contributes to the memory destabilization process (two-way ANOVA, F = 4.224, *P* = 0.0115; Bonferroni’s post-hoc test for within condition comparison and Newman-Keuls test for between conditions comparison; Fig. 1*C*,*D*). Ani administration alone resulted in almost complete fear memory amnesia of the weak AFC training (1FS-AFC), leaving no space for a further decrease in the tone-elicited freezing response. Therefore, autophagy induction combined with Ani (Ani+tBC) did not show any additional amnesic effect over Ani administration alone (two-way ANOVA, F (2, 22) = 1.594, *P* = 0.2257; Holm-Sidak’s post-hoc test; Fig. 1*C*,*E*).

### Autophagy overcomes a reconsolidation-resistant condition that is AMPAR endocytosis-dependent

Next, we examined the effect of autophagy induction on stronger AFC training by increasing memory strength using three tone-FS pairs (3FS-AFC), generating a reconsolidation-resistant condition. In the 3FS-AFC, Ani infusion into the LA after retrieval did not show any significant effect on the tone-elicited freezing response in comparison with the vehicle-infused condition. By contrast, Ani+tBC infusion after retrieval significantly reduced the tone-elicited freezing response levels, indicating that autophagy induction enhances memory destabilization beyond the fear memory reconsolidation-resistant condition. Without the retrieval session, Ani+tBC administration in the 3FS-AFC had no effect on auditory fear memory, indicating that a retrieval-specific process is necessary for the autophagy-enhancing effect on memory destabilization (two-way ANOVA, F (3, 30) = 3.476, *P* = 0.0281; Bonferroni’s post-hoc test for within condition comparison and -Kramer test for between conditions comparison; Fig. 2*A*,*B*). Collectively, the results obtained from both the inhibition and the induction of autophagy indicate a causal relationship between autophagy activity and memory destabilization.

**Figure 2.**
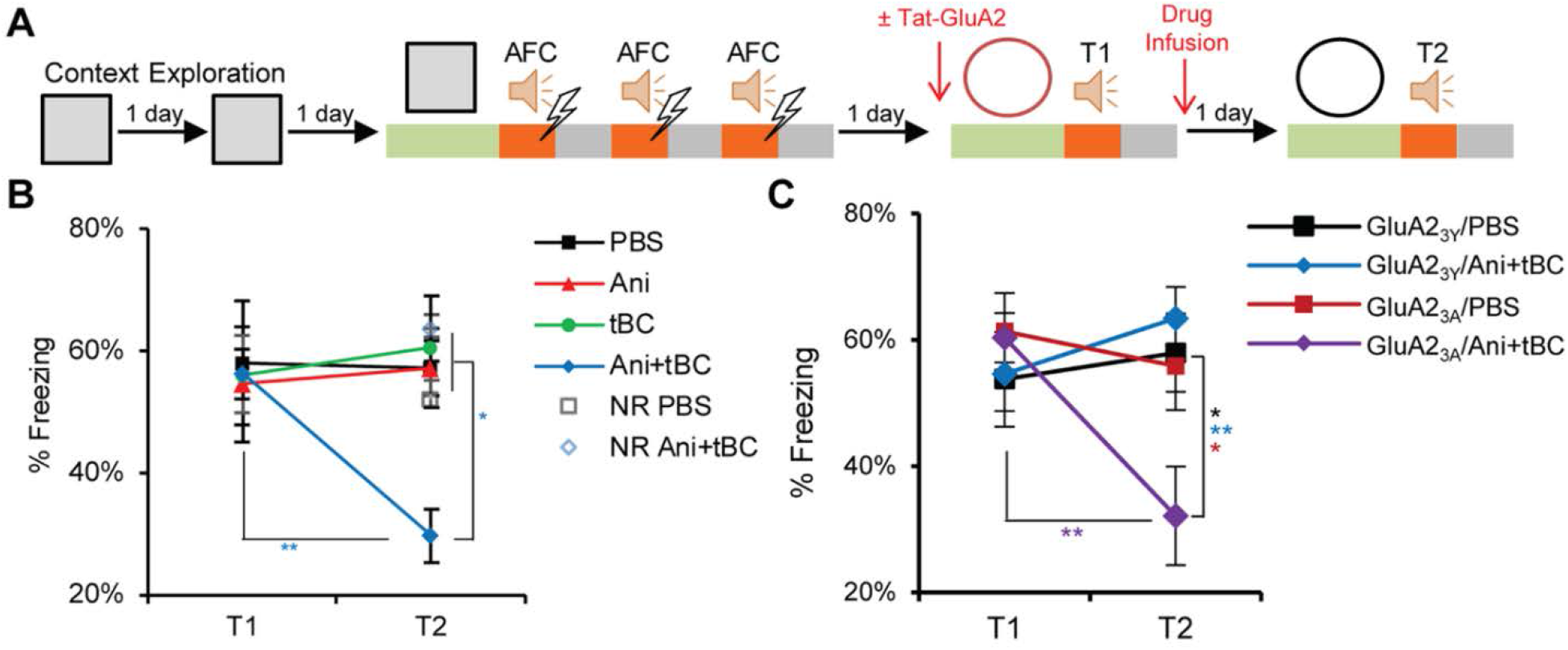
Autophagy overcomes a reconsolidation-resistant condition in an AMPAR endocytosis-dependent manner. ***A***, Design for the three tone-FS pairs auditory fear conditioning (3FS-AFC) experiments. The experiment was carried out either with no injection before T1 or with injection of Tat-GluA2 peptides: GluA2_3Y_, for blocking AMPA receptor endocytosis, or GluA2_3A_, as a negative control. ***B***, Average percentage freezing during tone at T1 and T2 showing that Ani combined with autophagy induction (Ani+tBC) showed significant retrograde amnesia while the Ani alone condition showed no amnesic effect (n = 7-10 mice/condition). ***C***, Average percentage freezing during tone at T1 and T2 showing that blocking AMPAR endocytosis abolished the amnesic effect of autophagy induction (n = 8-11 mice/condition). Error bars represent mean ± SEM; * *P* < 0.05; ** *P* < 0.01. Ani: anisomycin; NR: no reactivation (no T1); PBS: phosphate buffered saline; tBC: Tat-beclin 1.

We attempted to elucidate how autophagy modulates memory destabilization? As AMPAR are endocytosed after memory retrieval, we hypothesized that the autophagosome may fuse with endosomes carrying AMPAR and dictate their fate to lysosomal degradation (Rao-Ruiz et al., 2011; Shehata et al., 2012; Shehata and Inokuchi, 2014). Therefore, blocking endocytosis would block the autophagy effect on memory destabilization. The neural activity-dependent endocytosis of AMPAR relies on the carboxy-tail of GluA2, and the use of the synthetic peptide Tat-GluA2_3Y_ is well-established in attenuating activity-induced, but not constitutive, GluA2-dependent synaptic removal of AMPARs (Kim et al., 2001; Lee et al., 2002; Ahmadian et al., 2004; Scholz et al., 2010). In the 3FS-AFC, Tat-GluA2_3Y_ peptide infusion into the LA before retrieval completely blocked the Ani+tBC amnesic effect, while the control mutant peptide Tat-GluA2_3A_ had no effect (two-way ANOVA, F (3, 35) = 4.787, *P* = 0.0067; Bonferroni’s post-hoc test for within condition comparison and Tukey-Kramer test for between conditions comparison; Fig. 2*A*,*C*). These data indicate that AMPAR endocytosis is upstream to the autophagy induction effect on memory destabilization enhancement.

### Autophagy enhances retrograde amnesia of fear memory in CFC when targeted to the amygdala

We further investigated the autophagy induction effect on CFC as another reconsolidation paradigm. In CFC, the CS is a specific context, and the memory of the details of that context triggers a freezing response that is greater than that triggered by any other distinct context (Fanselow, 2000). Typically, inhibition of protein synthesis after CS retrieval leads to a certain degree of retrograde amnesia (Besnard et al., 2012; Finnie and Nader, 2012). To assess the degree of the retrograde amnesia, we compared it with that of a reference condition exposed to the same contexts without receiving any shock (NoFS). After CFC, an Ani infusion into the baso-lateral amygdala (BLA) after memory retrieval led to a decrease in the freezing response in comparison with the vehicle-infused condition (Fig. 3*A*,*B*) (Suzuki et al., 2004; Mamiya et al., 2009). Nevertheless, the freezing response after Ani administration was significantly higher than that in the NoFS condition, in both the specific and distinct contexts, implying that the resultant retrograde amnesia was only partial. After Ani+tBC administration, the average freezing response dramatically reduced, reaching no statistical significant difference from the NoFS condition in both contexts (two-way ANOVA, F (8, 102) = 10.19, *P* = 0.0001; Bonferroni’s post-hoc test for within condition comparison and Tukey-Kramer test for between conditions comparison; Fig. 3*B*). In addition, we assessed the complete amnesia for each mouse. In the Ani+tBC administered condition, 5 out of 12 mice were regarded as completely amnesic, in contrast, only 1 out of 12 mice in the Ani administered condition (see Materials & Methods section details) (Fig. 3 *C*,*D*). When the completely amnesic mice were subjected to a reconditioning session, they regained the freezing response to levels matching the pre-amnesic freezing levels, indicating an intact capacity for fear expression (one-way ANOVA, T1 and T2: *P* = 0.0001; within Ani+tBC complete group: *P* = 0.0021; Tukey’s post-hoc test; Fig. 3*E*). Altogether, these behavioral data indicate that induction of autophagy enhanced the amnesic effect of protein synthesis inhibition after retrieval and resulted in an enhanced level of retrograde amnesia.

**Figure 3.**
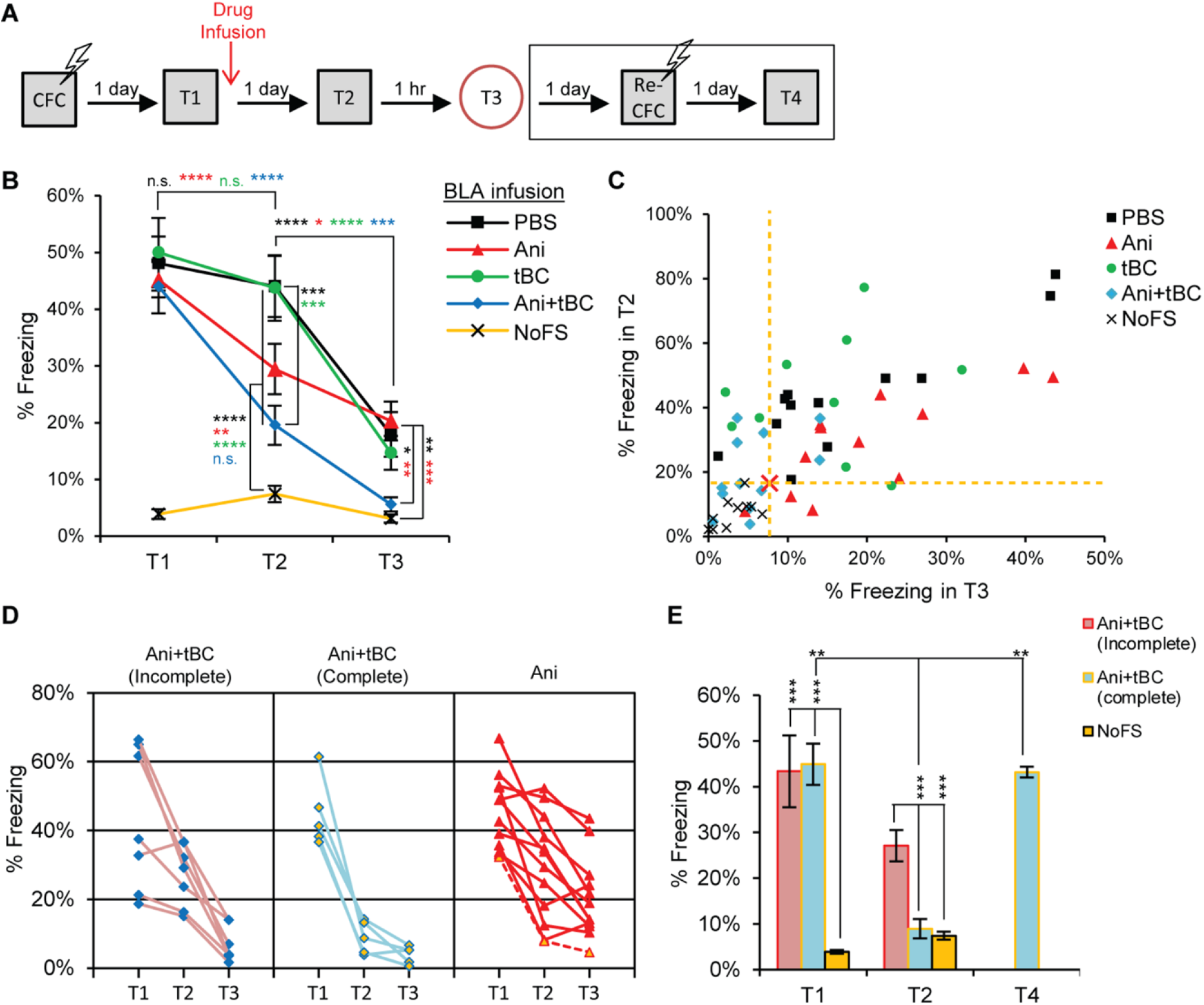
Autophagy enhances fear memory destabilization in contextual fear conditioning (CFC) when targeted to amygdala. ***A***, Design for the CFC reconsolidation and reconditioning experiments. ***B***, Average percentage freezing during retrieval (T1), and after the drugs were infused into the BLA when tested in the conditional stimulus context (T2) and in a distinct context (T3). Freezing levels at both T2 and T3 is showing a significant enhancement of anisomycin (Ani) amnesic effect on fear memory when combined with autophagy induction (n = 10-12 mice/condition). ***C***, Plot of individual mice freezing level at T2 against their freezing level at T3, for the assessment of complete fear amnesia after CFC. The red cross and yellow dashed lines represent a hypothetical point calculated from double the standard deviation for the freezing of the no foot shock (NoFS) condition at T2 and T3, where most of mice received Ani combined with Tat-beclin (Ani+tBC) treatment behaved as the NoFS condition. ***D***, Individual data for the complete and incomplete amnesic mice of the Ani+tBC condition compared with the Ani alone condition. The dashed red line is a complete amnesic mouse in the Ani only condition. ***E***, Average percentage freezing for the complete and incomplete amnesic mice of the Ani+tBC condition compared with the NoFS condition. The complete amnesic mice showed a normal freezing response one day after a reconditioning session (T4) (n = 5-10 mice/condition). Error bars represent mean ± SEM; * *P* < 0.05; ** *P* < 0.01; *** *P* < 0.001; **** *P* < 0.0001; n.s. = not significant. BLA: baso-lateral amygdala; PBS: phosphate buffered saline.

### Autophagy enhances retrograde amnesia of contextual memory when targeted to the hippocampus and AMPAR degradation in the spines of the memory-ensemble cells

We next examined the generality of the autophagy induction effect on other brain areas by targeting the CA1 region of the hippocampus. The tBC peptide induced autophagy in the hippocampus of adult mice when directly infused (one-way ANOVA, LC3-II/LC3-I: *P* = 0.0253 and Total LC3: *P* = 0.88; Tukey’s post-hoc test; Fig. 4*A*,*B*). As with the results from the BLA, Ani infusion into the CA1 after memory retrieval led to a decrease in the freezing response (two-way ANOVA, F (8, 106) = 9.027, *P* = 0.0001; Bonferroni’s post-hoc test for within condition comparison and Tukey-Kramer test for between conditions comparison; Fig. 4*C*,*D*) (Mamiya et al., 2009). As the hippocampal CA1 region encodes mainly spatial and contextual (CS) information, while the fear (FS or US) memory itself is encoded by the amygdala (LeDoux, 2000; Maren et al., 2013), Ani+tBC infusion into CA1 significantly decreased the discrimination between the specific and distinct contexts (two-way ANOVA, F (3, 44) = 7.211, *P* = 0.0005; Holm-Sidak’s post-hoc test for within condition comparison and Newman-Keuls test for between conditions comparison; Fig. 4*E*,*F*) without affecting the fear memory itself in the CS context (Fig. 4*D*,*E*). The same result was obtained when another autophagy inducer, trifluoperazine (TFP), was combined with Ani (two-way ANOVA, F (3, 18) = 5.328, *P* = 0.0084; Bonferroni’s post-hoc test for within condition comparison and Tukey-Kramer test for between conditions comparison; Fig. 4*G*). These results indicate that the enhanced memory destabilization resulting from induction of autophagy is not restricted to one brain area, and that the behavioral outcome of autophagy induction differs in accordance with the main information encoded in the target brain area.

**Figure 4.**
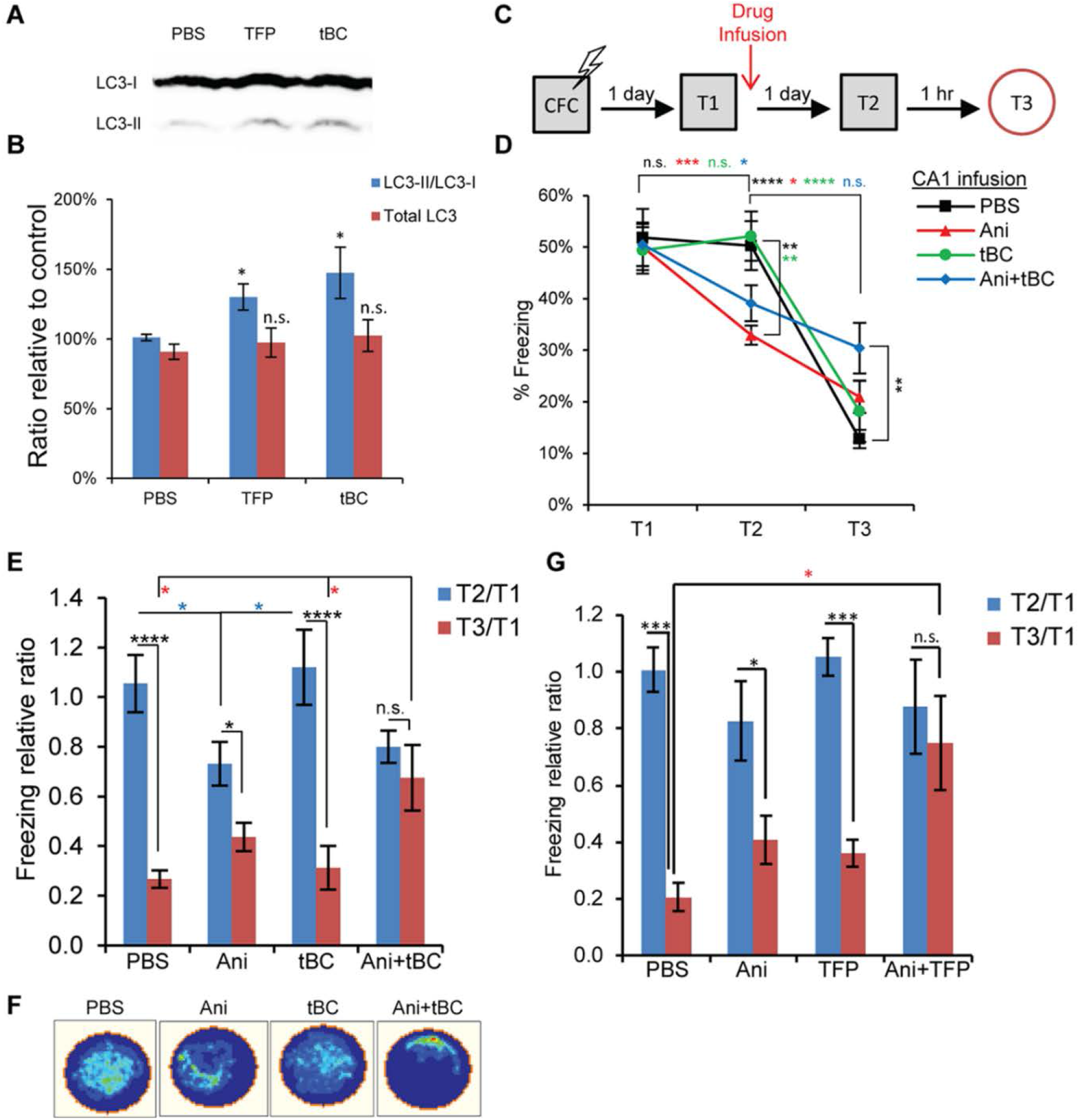
Autophagy enhances context memory destabilization in contextual fear conditioning (CFC) when targeted to hippocampus. ***A***, Representative immunoblot from hippocampal lysates collected 4 h after unilateral drug infusion into CA1. ***B***, Quantitation of the signal intensity represented as percentage relative to a control PBS sample (n = 4-5 per condition). ***C***, Design for the CFC reconsolidation experiment. ***D***, Average percentage freezing during retrieval (T1), and after the drugs were infused into the hippocampal CA1 region when tested in the conditional stimulus context (T2) and in a distinct context (T3). Combined anisomycin and autophagy induction (Ani+tBC) showed higher freezing levels at T3 compared to Ani alone condition (n = 10-14 mice/condition). ***E***, Data from ***D*** represented as the freezing level at T2 or T3 relative to T1, showing the loss of context discrimination after Ani+tBC combined treatment. ***F***, A representative occupancy plot for a mouse per condition at T3 from D. ***G***, Freezing level at T2 or T3 relative to T1 after autophagy induction by infusion of TFP into the hippocampal CA1 region alone or combined with Ani. As with tBC, TFP combined with Ani resulted in loss of context discrimination (n = 5-6 mice/condition). Error bars represent mean ± SEM; * *P* < 0.05; ** *P* < 0.01; *** *P* < 0.001; **** *P* < 0.0001; n.s. = not significant. PBS: phosphate buffered saline.

We tested the involvement of autophagy in the degradation of the endocytosed AMPAR after retrieval utilizing the CFC model and benefiting from the dendrite orientation in the CA1 radiatum. We quantified the level of AMPAR co-localizing with the spines of the neurons holding the memory trace after the amnesic treatments. To label the memory-ensemble cells in the CA1 region, lentivirus expressing mCherry under the control of the tetracycline response element was injected into c-fos-tTA transgenic mice, which had been maintained on a diet containing doxycycline, except for the period spanning 2 days before and 6 h after the CFC session (Fig. 5*A*,*B*) (Reijmers et al., 2007; Ohkawa et al., 2015). Following retrieval, vehicle or Ani+tBC was unilaterally infused into the CA1, and changes in the GluA1, an AMPAR subunit, staining and its overlap with the mCherry spines (representing the spines of the memory-ensemble cells) were checked 1 day later, reflecting their status at the test 2 session (Fig. 5*B*,*C*). The GluA1 puncta were classified into strong or weak signals according to their fluorescent intensity, and reflecting the level of AMPAR enrichment. The total GluA1 puncta, the ratio of strong signals to the total GluA1 puncta, and the number of mCherry-only spines did not significantly differ between the two conditions (unpaired Student’s t-tests; Fig. 5*D*,*E*, *I-K*). Only the overlap between the GluA1 strong signals and mCherry spines was significantly lower in the Ani+tBC condition compared to the vehicle condition and not significantly different from the chance level (two-way ANOVA, F (1, 4) = 12.4, *P* = 0.0244; Bonferroni’s post-hoc test; Fig. 5*F*-*H*). This decrease in AMPAR enrichment in the spines of ensemble-cells to the chance level is in accordance with the behavioral data showing a decrease in contextual memory and hence loss of context discrimination (Fig. 4*E*).

**Figure 5.**
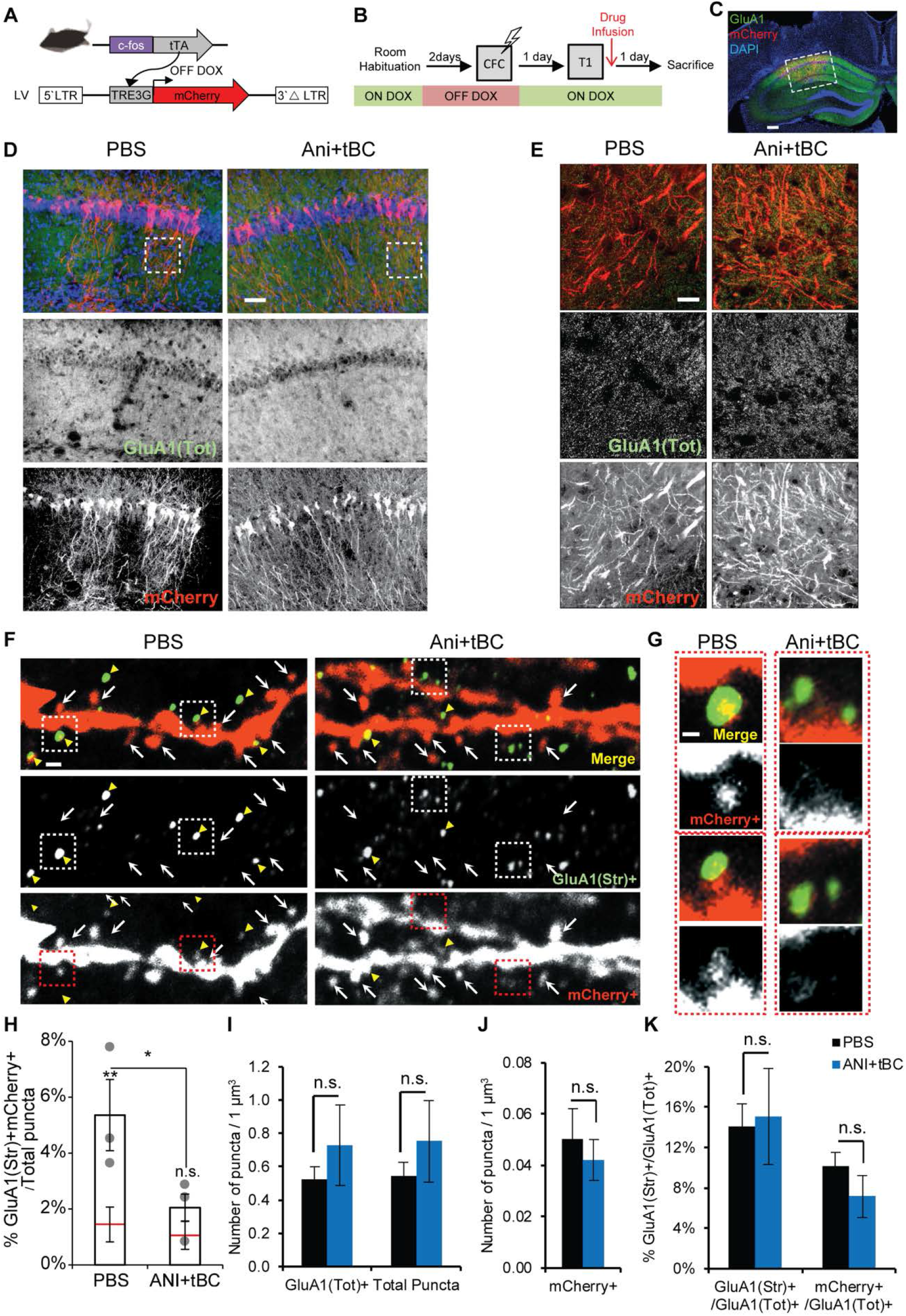
Autophagy enhances AMPA receptor degradation in the spines of memory-ensemble cells. ***A***, Lentivirus-mediated labeling of the spines of the memory-ensemble cells with mCherry in the c-fos-tTA transgenic mice. ***B***, Experimental design for checking the effect of autophagy induction on α-amino-3-hydroxy-5-methyl4-isoxazolepropionic acid receptor (AMPAR) expression and distribution in memory-ensemble cells. ***C***, Images showing immunohistochemical staining for mCherry (red), and endogenous GluA1 (green) in an Ani+tBC treated hemisphere (scale bar, 400 μm). ***D-E***, Low magnification images (D; scale bar, 50 μm) and higher magnification maximum intensity projection images (E; scale bar, 20 μm) showing that anisomycin combined with Tat-beclin (Ani+tBC) treatment did not affect the overall AMPA receptor signals compared to PBS control. ***F-G***, Representative dendrites for each treatment condition showing less co-localization of the mCherry-stained spines (mCherry^+^) with the GluA1-strongly-stained puncta (GluA1(Str)^+^) in the Ani+tBC condition than in the PBS condition (scale bar, 500 nm; insets are shown in G; co-localization is indicated by yellow arrowheads, and arrows indicate mCherry^+^-only spines). ***G***, Higher magnification images for two spines per condition (scale bar, 200 nm). ***H***, Quantitation for the co-localization of mCherry^+^ spines with the GluA1(Str)^+^ puncta per total puncta counted and the chance level (red line). The overlap between mCherry^+^ spines and the GluA1(Str)^+^ decreased to chance level after Ani+tBC treatment (n = 3 hemispheres/condition; four images/hemisphere). ***I-K***, No significant difference between PBS- or Ani+tBC-injected hemispheres in: ***I***, The total GluA1 puncta (GluA1(Tot) ^+^), or total counted puncta. ***J***, The mCherry-labeled spines (mCherry^+^). ***K***, The ratio of the GluA1 puncta with a strong signal (GluA1(Str) ^+^) or in the mCherry^+^ spines with GluA1(Tot) ^+^ puncta (n = 3 hemispheres/condition; four images/hemisphere). Error bars represent mean ± SEM; * *P* < 0.05;; n.s. = not significant. DOX: doxycycline; PBS: phosphate buffered saline.

### Autophagy destabilizes synaptic plasticity in a LTP reconsolidation model

Finally, we tested the effect of autophagy induction on synaptic destabilization using an *in vivo* LTP system in rats, in which a protein synthesis-dependent long-lasting LTP was induced in the dentate gyrus by 400 Hz high frequency stimulation of the perforant path (unpaired Student’s t-test, 400 Hz: *P* = 0.0371; Fig. 6*A*,*B*) (Fukazawa et al., 2003). To model synaptic reconsolidation, the perforant path was reactivated by a protein synthesis-dependent 8 Hz stimulation (unpaired Student’s t-test, 8 Hz: *P* = 0.0086; Fig. 6*B*) 1 day after LTP induction; this to resensitize the LTP to the protein synthesis inhibitor Ani, thereby mimicking behavioral reconsolidation (Fig. 6*C*) (Okubo-Suzuki et al., 2016). Ani treatment significantly decreased the field excitatory postsynaptic potential (fEPSP) slope 1 day after 8 Hz reactivation compared with the vehicle condition. However, this effect was only partial, as the fEPSP slope was still higher than the baseline level before LTP induction. Following Ani+tBC treatment, LTP destabilization was almost complete and the fEPSP slope was not significantly higher than the baseline level. This enhancement of synaptic destabilization was not observed when Ani administration was combined with the unfused Tat peptide (D-tat) (two-way ANOVA, F (20, 235) = 4.279, *P* = 0.0001; Holm-Sidak’s post-hoc test; Fig. 6*D*,*E*). These data indicate that induction of autophagy enhanced the synaptic destabilization triggered by the 8 Hz reactivation. Furthermore, behavioral reconsolidation is dependent on NMDA receptors (NMDAR), and GluN2B-containing NMDAR (NMDAR-2B) is required for memory destabilization after recall (Ben Mamou et al., 2006; Milton et al., 2013). In our synaptic reconsolidation model, ifenprodil, a selective NMDAR-2B antagonist, blocked the LTP-destabilizing effect of Ani administration, mimicking behavioral reconsolidation. Also, ifenprodil completely blocked the LTP-destabilizing effect of Ani+tBC administration (two-way ANOVA, F (10, 105) = 0.5138, *P* = 0.8771; Holm-Sidak’s post-hoc test; Fig. 6*F*,*G*). These data indicate that NMDAR-2B was involved in physiological destabilization in our *in vivo* synaptic reconsolidation model, and demonstrate that the effect of enhanced autophagy on synapse destabilization is downstream of NMDAR-2B.

**Figure 6.**
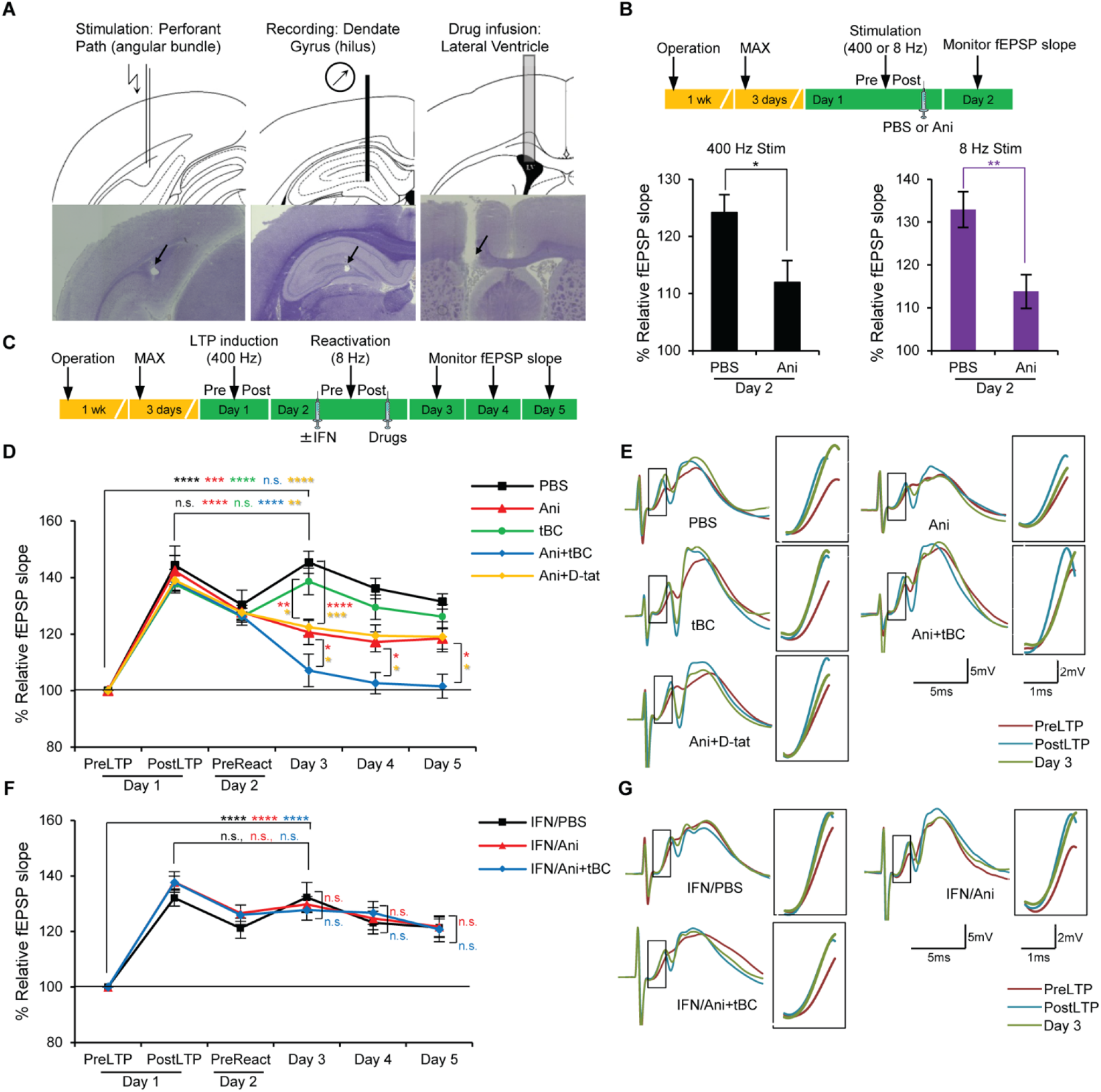
Autophagy induction enhances synaptic destabilization of long-term potentiation (LTP) in freely moving rats. ***A***, Diagrams and images of hematoxylin-stained slices for the stimulation electrode, the recording electrode, and the drug injection cannula. Arrows indicate corresponding scars. ***B***, In the *in vivo* LTP, both 400 and 8 Hz stimulation were protein synthesis-dependent. Anisomycin (Ani) or phosphate buffered saline (PBS) was infused 5 minutes after the 400 or 8 Hz stimulation and percentage field excitatory postsynaptic potential (fEPSP) slope was calculated on day 2 relative to the pre-stimulation level on day 1 (n = 5-7 rats/condition). ***C***, Design for the LTP reconsolidation experiment; fEPSP was recorded immediately before (pre) and after (post) LTP induction and reactivation, and for 3 consecutive days after intracerebroventricular drug infusion. ***D***, Percentage fEPSP slope relative to the preLTP level showing that autophagy induction using Tat-beclin 1 (tBC), but not the unfused control peptide D-tat, significantly enhanced synaptic destabilization compared to anisomycin (Ani) only treatment. Note: no injections were given before reactivation (n = 10-11 rat/condition). ***E***, Representative waveform traces and enlarged portion of slope (inset) for each treatment from D. ***F***, Percentage fEPSP slope relative to the preLTP level when ifenprodil (IFN), a N-methyl-D-aspartate (NMDA) receptor blocker, was injected before 8 Hz reactivation. IFN completely blocked the synaptic destabilization effect of the Ani only and the Ani+tBC treatments (n = 8 rats/condition). *G*, Representative waveform traces and enlarged portion of slope (inset) for each treatment from F. Error bars represent mean ± SEM; * *P* < 0.05; ** *P* < 0.01; *** *P* < 0.001; **** *P* < 0.0001; n.s. = not significant.

## Discussion

Our results indicate that autophagy contributes to memory destabilization and that its induction enhances memory destabilization, including a reconsolidation-resistant one, and the degradation of the endocytosed AMPAR in the spines of memory ensemble-neurons. Also, autophagy induction enhances synaptic destabilization in an NMDAR-dependent manner (Fig. 7*A*,*B*). A consistent finding through our study is that autophagy induction alone, through tBC administration after reactivation, did not show any significant amnesic effect in the 3FS-AFC, CFC, and synaptic reconsolidation models. This indicates that, regardless of the degree of destabilization, if the protein synthesis is not compromised within a certain time window following reactivation, the synthesized proteins have the capacity to regain synaptic plasticity and memory (Fig. 7*B*). This demonstrates the capacity of protein synthesis in the restabilization of synapses and the reinstating of specific memories.

**Figure 7.**
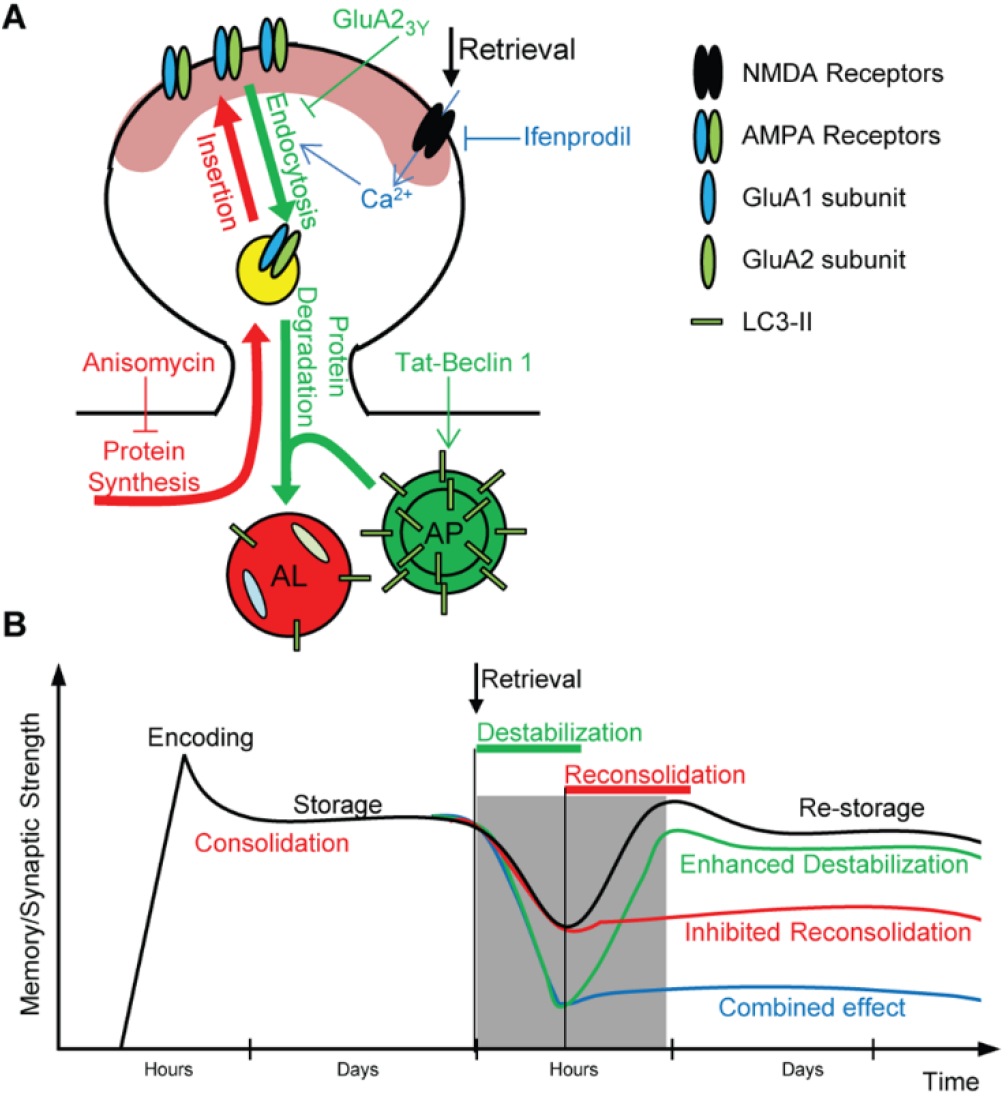
Models for the effect of autophagy on synaptic and memory destabilization. ***A***, Molecular mechanism for the effect of autophagy induction on synaptic and memory destabilization. Memory retrieval leads to NMDAR activation, which stimulates both autophagy and the endocytosis of AMPAR in the activated neurons. Autophagosomes (AP) fuse with the endosomes carrying internalized AMPARs, forming autolysosomes (AL), and dictate their fate to lysosomal degradation (green arrows). During the reconsolidation process, newly synthesized proteins, including AMPARs, are delivered to the synaptic surface replacing the degraded ones (red arrows). In the present study, ifenprodil (IFN) was used to block NMDA receptor activation, GluA2_3Y_ peptide to block endocytosis of AMPARs, Tat-beclin-1 (tBC) peptide to induce autophagy, and anisomycin (Ani) to inhibit protein synthesis. ***B***, Model explaining the time line of synaptic and memory strength changes by autophagy. After recall, consolidated synaptic plasticity and memory strength are usually physiologically destabilized and return to a labile state, after which a protein synthesis-dependent reconsolidation process is required for restabilization (black line). The labile state (destabilization) is inferred from the decreased synaptic strength and the retrograde amnesia produced by the protein synthesis inhibition (red line). Autophagy induction when combined with protein synthesis inhibition leads to a greater decrease in synaptic strength and enhanced retrograde amnesia, indicating enhanced destabilization (blue line). Autophagy induction alone does not affect synaptic or memory strength (green line).

AMPAR are heterotetrameric complexes composed of various combinations of four subunits (GluA1−4), with the GluA1/2 and GluA2/3 tetramers being the two major subtypes (Wenthold et al., 1996). The amount of synaptic GluA2-containing AMPARs correlates with LTM maintenance and strength (Yao et al., 2008; Migues et al., 2010; Migues et al., 2014; Dong et al., 2015). Blocking the endocytosis of GluA2-containing AMPARs inhibits the induction of LTD, but not LTP, without affecting basal synaptic transmission (Ahmadian et al., 2004; Brebner et al., 2005; Dalton et al., 2008; Scholz et al., 2010). More relevant is its involvement in memory destabilization, where it does not affect the acquisition or the retrieval of conditioned fear memory (Rao-Ruiz et al., 2011; Hong et al., 2013). These reports are in agreement with our hypothesis that autophagy works through dragging endosomes carrying AMPAR to lysosomal degradation, as evidenced by our demonstration that GluA2-dependent AMPAR endocytosis is a prerequisite for autophagy to affect memory destabilization. Additionally, GluA2-dependent AMPAR endocytosis correlates with the decay of LTP and the natural active forgetting of LTM (Hardt et al., 2014; Dong et al., 2015; Migues et al., 2016), which suggests that autophagy may play a role in the forgetting of consolidated memories through the gradual synaptic loss of AMPAR overtime, and, hence, memory loss.

The GluA1 subunit acts dominantly over other subunits to determine the direction of AMPAR to the surface, and is correlated with synaptic potentiation, LTP, and fear memory (Ehlers, 2000; Shi et al., 2001; Malinow and Malenka, 2002; Lee et al., 2004; Rumpel et al., 2005). A mouse model lacking GluA1 subunit expression exhibits impaired hippocampus-dependent spatial memory (Reisel et al., 2002; Sanderson et al., 2007). CFC recruits newly synthesized GluA1-containing AMPAR into the spines of the hippocampal memory-ensemble cells in a learning-specific manner (Matsuo et al., 2008). Inhibitory avoidance, a hippocampus-dependent contextual fear-learning task, delivers GluA1-containing AMPARs into the CA1 synapses in the dorsal hippocampus, where they are required for encoding contextual fear memories (Mitsushima et al., 2011). Inhibition of the cAMP response element-binding protein, a key transcription factor implicated in synaptic plasticity and memory, is associated with a specific reduction in the AMPAR subunit of GluA1 within the postsynaptic densities, and impaired CFC (Middei et al., 2013). These reports are in agreement with our use of spine enrichment with GluA1-containing AMPAR as a molecular reflection of synaptic and contextual memory strength in the CA1 region of the hippocampus.

D-cycloserine, an NMDAR agonist, prepares resistant memories for destabilization (Bustos et al., 2010). Noteworthy, d-cycloserine also enhances memory update (fear extinction) by increasing GluA2-containing AMPAR endocytosis, and augments NMDAR-2B-dependent hippocampal LTD (Duffy et al., 2008; Bai et al., 2014). Therefore, autophagy might be a potential downstream mechanism by which d-cycloserine facilitates destabilization.

In the present study using the CFC paradigm, manipulation of BLA neurons led to complete retrograde amnesia, while manipulation of CA1 neurons led to a generalization of fear. Therefore, targeting of the proper brain region is necessary to achieve the desired behavioral response. This highlights the importance of targeting the fear memory to successfully alleviate PTSD symptoms, rather than any other associated memory within the entire network.

We showed here that autophagy destabilizes resistant memories formed under stressful conditions, suggesting autophagy as a potential target for clinical applications. Owing to the growing interest in finding autophagy inducers for several applications, many FDA-approved autophagy inducers already exist, including known anti-psychotic and anti-depressant drugs, and more specific ones are on their way (Levine et al., 2015; Morel et al., 2017). This increases the feasibility of using autophagy inducers for future therapeutic applications, including PTSD treatment.

## Acknowledgements

This work was supported by the Grant-in-Aid for Scientific Research on Innovative Areas ‘Memory dynamism’ (JP25115002) from the Ministry of Education, Culture, Sports, Science, and Technology (MEXT), JSPS KAKENHI grant number JP23220009, the Core Research for Evolutional Science and Technology (CREST) program (JPMJCR13W1) of the Japan Science and Technology Agency (JST), the Mitsubishi Foundation, the Uehara Memorial Foundation, and the Takeda Science Foundation support to K.I.; and by a Grant-in-Aid for young scientists from JSPS KAKENHI grant number JP25830007 to M.S. The Otsuka Toshimi Scholarship Foundation supported K.A.

M.S. and K.I. designed the experiments and wrote the manuscript. M.S., K.C., and K.A. performed the experiments. M.S., K.A., K.C., and K.I. analyzed the data. H.N. and M.M. produced and maintained transgenic mice. M.S. and K.A. contributed equally to this work.

From the University of Toyama, we thank N. Ohkawa for assistance with c-fos-tTA mice, R. Okubo-Suzuki and Y. Saitoh for assistance with electrophysiology, S. Kosugi for lentivirus preparation, and S. Tsujimura for maintenance of mice.

